# The dendritic spine vicinome: characterizing its crowding in the human brain with petavoxel electron microscopy

**DOI:** 10.64898/2026.09.16.752196

**Authors:** Elias Manjarrez, Nicte Lozada, Miguel A. Zamora-Ursulo, Amira Flores

## Abstract

For more than a century we have pictured the dendritic spine as the postsynaptic point where a single presynaptic bouton delivers its message. That picture is correct but incomplete, since it overlooks other neuronal elements surrounding each spine. Here we characterize that neighborhood for the first time by counting and measuring the proximity of the elements encircling each postsynaptic spine. We call this neighborhood the vicinome: the full set of neuronal and glial elements surrounding a dendritic spine. We used ground-truth three-dimensional meshes from the H01 petavoxel dataset, a reconstruction of the human cerebral cortex. We analyzed 4550 neighboring elements across 322 spines from 17 pyramidal neurons in cortical layers 2 through 5, with each spine classified as apical or basal. Neighbor number followed a depth gradient, falling from 17.2 ± 4.2 elements per spine in layer 2 apical spines to 12.5 ± 3.1 in layer 5 basal spines. A linear mixed model with a random intercept per neuron confirmed a strong effect of layer (p = 4.3×10^-5^) with large effect sizes (Cohen’s d up to 1.45) and no effect of polarity. The minimum spine-to-neighbor distance stayed uniform, near 30 nm in every group, and although some depth-related contrasts were significant, all effect sizes were negligible (Cohen’s d below 0.21). We conclude that cortical depth, and not pyramidal neuron polarity, was related to the number of neuronal elements neighboring the spine. We suggest that vicinome crowding could offer a measurable observable for comparison across regions, ages, species, and disease.

**New & Noteworthy:** Using the H01 petavoxel reconstruction of the human cortex, we characterize, for the first time, the neuronal neighborhood of the dendritic spine we term the dendritic spine vicinome. Cortical depth, not dendritic polarity, governs how many neuronal elements crowd around a spine. Yet those neighbors stay uniformly close across all layers. The spine is not an isolated synaptic point, but a vicinome hub embedded in a dense envelope.

## Introduction

The dendritic spine has been the emblem of the postsynaptic compartment since Ramón y Cajal first described the dendritic surface as studded with short spines on Purkinje cells (García-López et al., 2007, 2010; Yuste, 2015). For more than a century, the spine was understood mainly as the receiving end of a single excitatory contact, the point where one presynaptic bouton meets one postsynaptic density across a cleft of a few tens of nanometers (Tønnesen and Nägerl, 2016; Südhof, 2018; Caire et al., 2023). However, in 2019, Kwon et al. found that a spine can be activated by neurotransmitter released from a neighbor that never contacts it across a classical synapse. The synaptic influence travels through the space between membranes to activate extrasynaptic receptors. This finding suggested that the dendritic spine does not sit in empty space with a single synaptic contact. It is embedded in dense neuropil, surrounded by boutons, dendrites, spines, and glial processes from many other neurons packed within nanometers of its membrane that may influence it, supporting neuronal ensembles in cortical circuits (Geckt et al., 2026). However, how crowded this immediate neighborhood is, and how close these elements lie, has remained largely unmeasured.

Researchers have scrutinized the spine’s form for decades. Its head and neck vary along a continuum of shapes that shifts with age and species (Ofer et al., 2022), and three-dimensional reconstructions from Golgi impregnation to serial electron microscopy have mapped that diversity in the human brain with growing precision (Benavides-Piccione et al., 2013; Reberger et al., 2018; Renner and Rasia-Filho, 2023). The geometry is not idle, since the size and shape of a spine track the strength and stability of the synapse it carries (Arellano et al., 2007; Bourne and Harris, 2008), and the neck in particular tunes how the signal born at the head reaches the dendrite (Wilson et al., 1983; Harris and Stevens, 1989; Rochefort and Konnerth, 2012). A century of work has told us, in fine detail, what a spine looks like and how its shape shapes its function. What it has left largely unaddressed is the surrounding neighborhood.

In support of that crowded neighborhood, Hwang et al. (2021) employed three-dimensional electron microscopy and genetically identified neurons in the mouse primary visual cortex to show that the perisomatic membrane of distinct GABAergic cells organizes excitatory and inhibitory inputs in a cell-type-specific way, and that a single spine on these interneurons can carry as many as eight synaptic inputs rather than one. More recently, Glausier et al. (2025) applied the same volume approach to postmortem human dorsolateral prefrontal cortex, densely reconstructing the neuropil around individual synapses and recovering intact ultrastructural correlates of synaptic function, showing that the crowded environment of the human dendritic spine is now open to quantitative three-dimensional analysis. However, in their study the neuropil was reconstructed to characterize the synapse and its subcellular machinery, not to ask how many elements encircle a given spine or how near they lie. The neighborhood itself, its crowding and its geometry, was never the unit of measurement.

The recent release of the H01 dataset, a cubic millimeter of human cerebral cortex from the anterior temporal lobe reconstructed at nanometer resolution and navigable through Neuroglancer (Shapson-Coe et al., 2024; Januszewski et al., 2018), removes the last barrier and makes the full neuronal neighborhood of a human dendritic spine accessible to direct measurement. This H01 dataset was obtained using automated tape-collecting ultramicrotomy (ATUM) with scanning EM (SEM) (Kubota et al., 2018).

Neighborhood crowding matters because spines integrate signals. A spine’s output in response to one or several synaptic inputs depends not only on the input it receives but also on its local electrical and biochemical environment, which surrounding elements help shape. Neighboring membranes constrain the extracellular space, and the extracellular fields generated by active neurons can feed back onto the membrane potential of nearby cells independently of chemical or electrical synapses, a phenomenon known as ephaptic coupling that is strongest where elements lie close together and at slow timescales (Anastassiou et al., 2011; Han et al., 2018). A spine encircled by many tightly apposed neighbors therefore inhabits a different integrative context than an isolated one. Electron microscopy alone cannot establish whether such field interactions occur, but it can measure the anatomical proximity that any of these interactions, chemical, electrical, or ephaptic, would require, and proximity is the necessary substrate on which they rest.

Here we characterize, for the first time, the neuronal neighborhood of the human dendritic spine. Navigating the H01 reconstruction in Neuroglancer, we manually identified 4550 neighboring neuronal elements surrounding 322 spines from 17 pyramidal neurons across cortical layers 2 through 5, and from the ground-truth three-dimensional meshes we quantified two properties of each neighborhood: the number of neighboring elements per spine and the minimum surface-to-surface distance between the spine and those elements. We further asked whether these properties differ with cortical depth and with the apical or basal polarity of the parent dendrite. This characterization shifts the description of the spine from an isolated synaptic point to the crowded three-dimensional environment of the vicinome in which it operates, allowing us to measure the degree to which those elements press around the spine as an observable that can be quantified and compared. The vicinome concept matters because it is not the neuropil as a whole, but a delicate and organized portion of it that surrounds a single spine and that is measured relative to that spine.

## Materials and Methods

### Dataset

We based this study on H01, the openly released nanoscale connectomic volume of human cortex (Shapson-Coe et al., 2024). The specimen originated from a small block of anterior middle temporal gyrus removed during epilepsy surgery, spanning roughly one cubic millimeter of tissue. After heavy-metal staining, it was cut into serial sections of approximately 33 nm and captured with multibeam scanning electron microscopy at a lateral resolution of 4 by 4 nm. Volumetric alignment followed by flood-filling network segmentation (Januszewski et al., 2018) reconstructed nearly every neuron and process as a three-dimensional mesh, refined through conservative c3 agglomeration. From this reconstruction, we sampled 322 dendritic spines from 17 pyramidal neurons spanning cortical layers 2 through 5. Following Ramón y Cajal’s criteria for pyramidal dendritic arborization, we assigned each spine to the apical or basal compartment of its parent dendrite.

### Spine and neighbor identification in Neuroglancer

A trained observer identified spines and their surrounding neuronal elements in Neuroglancer, the browser-based viewer distributed with H01 and developed by J. Maitin-Shepard and the Google Connectomics team. Neuroglancer presents three linked panels. A two-dimensional panel displays any pair of the X, Y, and Z axes for voxel selection, a three-dimensional panel renders the selected object for free rotation, panning, and zooming, and a control panel selects cortical layer, cell type, and individual segments while recovering each object’s unique identifier for later analysis. For each spine, the observer inspected the segmented volume and recorded the identity of each neighboring element in physical contact with, or immediately surrounding, the spine, including presynaptic boutons, dendrites and spines of other neurons, and glial processes. This manual annotation established each spine’s neighbor set, which served as the ground truth for subsequent measurements.

### Ground truth mesh retrieval and measurement

We obtained both the number of neighbors and the spine-to-neighbor distances directly from the ground-truth three-dimensional meshes of the H01 reconstruction, consistent with the mesh-based morphometric approach validated in our preceding reports (Manjarrez et al., 2026; Zamora-Ursulo and Manjarrez, 2026). This ensures the present study rests on the same objective surface geometry rather than projected visual estimates.

We retrieved segmentation meshes from the publicly available c3 reconstruction at native voxel resolution (8 by 8 by 33 nm in x, y, and z) and processed them in Python. For each spine, we cropped its mesh and each annotated neighbor’s mesh to a three-micrometer spherical region of interest centered on the spine coordinates, converted from voxel units to nanometers. When cropping left fewer than 50 vertices on a mesh, we widened the sphere to 4.5 micrometers and excluded neighbors that still yielded fewer than 4 vertices. Raw meshes from the voxel grid carry staircase-like surface artifacts, so we applied Laplacian smoothing with 5 iterations and a relaxation factor of 0.1 before any distance computation. This step refines local surface geometry without distorting the structure’s overall morphology.

### Number of neighboring elements

For each spine, we counted the number of annotated neighbors that survived the mesh criteria above. This count defines the local crowding of the spine neighborhood. Because both the counts and the distances derive from the same cleaned population of neighbors, the number of neighbors counted per spine equals the number of spine-to-neighbor distances measured, which keeps the two analyses internally consistent.

### Minimum spine-to-neighbor distance

We computed the minimum membrane-to-membrane distance bidirectionally. For every vertex on the spine surface, we queried the nearest point on the neighbor surface, and we then repeated the procedure in the opposite direction. The smallest value across both passes was taken as the distance for that spine-to-neighbor pair. This two-way approach prevents underestimation when meshes differ substantially in vertex density, which is common when comparing small spine fragments against large neuronal or glial processes. We discarded no distances based on magnitude, preserving the full biological range for analysis.

### Statistical analysis

We organized data into eight groups defined by cortical layer (L2-L5) crossed with spine polarity (apical or basal). For each variable, the number of neighbors and the minimum distance of the vicinome, we first tested a global difference across the eight groups with the Kruskal-Wallis test. Because spines are nested within neurons and are therefore not independent, the primary analysis was a linear mixed model (LMM) with layer and polarity as fixed effects and a per-neuron random intercept to absorb baseline variability between individual neurons. We report the omnibus effects of layer, polarity, and their interaction. We used pairwise Mann-Whitney tests among the eight groups (all 28 combinations) with Benjamini-Hochberg false discovery rate (FDR) correction, and considered values below 0.05 significant. Because these groups are independent samples, we expressed effect sizes as independent-samples Cohen’s d with its 95% confidence interval. We presented data in violin plots showing individual values, group mean, and the standard error of the mean. All analyses were performed in MATLAB R2025a (The MathWorks, Inc., Natick, MA, USA) using the Statistics and Machine Learning Toolbox.

## Results

### Manual identification of the dendritic spine vicinome in Neuroglancer

To examine the neuronal elements surrounding each dendritic spine (the vicinome), we first manually navigated the H01 reconstruction in Neuroglancer, inspecting each spine along with the segmented processes around it. This manual step served two purposes. It revealed the composition of each neighborhood in three dimensions and fixed the spatial coordinates of every spine whose neighborhood we explored. We then passed those coordinates to our Python routine, which measured the minimum spine-to-neighbor distances on the ground-truth meshes with the precision reported in the following section.

We analyzed 4550 neighboring elements across 322 spines from 17 pyramidal neurons in cortical layers 2 through 5, with each spine classified as apical or basal. As representative examples, we show the neighborhoods of two typical dendritic spines, each surrounded by several neuronal elements. **Figure 1** shows a spine, colored green, belonging to a layer 5 pyramidal neuron (segment ID 3687613997). Panel A shows the spine alone, protruding from its parent dendrite. Panels B through H add the neighboring neuronal elements one at a time, so that the neighborhood of the dendritic spine vicinome is built up sequentially around the same spine. The first neighbor appears in yellow (B), followed by a light blue element (C), a purple one (D), a red one (E), a second purple element (F), a magenta one (G), and finally a light green element (H). The two purple elements in panels D and F belong to two different neighboring neurons and are not the same process, although Neuroglancer rendered them in the same color. By the last panel (H), the spine that stood alone in A is enclosed within a dense cluster of neuronal elements from neighboring neurons, illustrating the crowding of its neighborhood.

**Figure 1.**
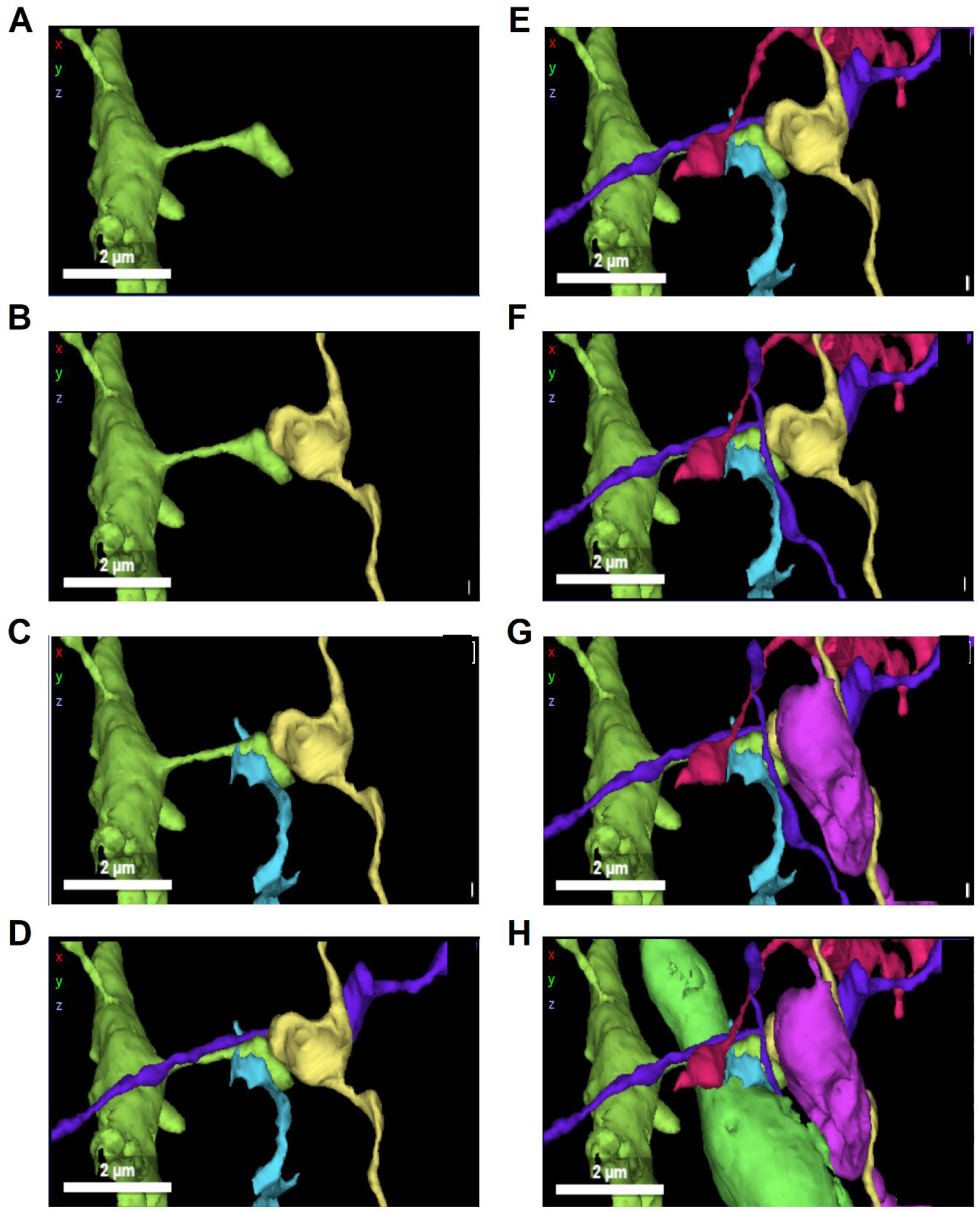
Illustrating the dendritic spine vicinome with an example. Sequential reconstruction of the neuronal neighborhood of a layer 5 dendritic spine. A dendritic spine (green) belonging to a layer 5 pyramidal neuron (segment ID 3687613997) is shown with the surrounding neuronal elements, retrieved from the H01 reconstruction and rendered in three dimensions in Neuroglancer. **(A)** The spine alone, protruding from its parent dendrite. **(B through H)** The neighboring elements are added one at a time to build up the neighborhood around the same spine, in the order yellow (B), light blue (C), purple (D), red (E), a second purple element (F), magenta (G), and light green (H). By panel H, the spine is enclosed within a dense cluster of processes from other neighboring neurons, illustrating the crowding of its neighborhood. The two purple elements in panels D and F belong to two different neighboring neurons and are not the same process; Neuroglancer assigned them the same color by coincidence. Likewise, the light green neighbor added in panel H is a separate element from the green spine and only shares a similar hue. Scale bars: 2 micrometers.

**Figure 2** shows a second spine and its vicinome, again colored green, from the same layer 5 neuron, this time shown with orthogonal electron microscopy sections through the spine head. As in the previous figure, panel A shows the spine alone, and panels B through H add the surrounding neuronal elements in sequence, shown in both the three-dimensional rendering and the corresponding cross-sectional planes. The orthogonal views confirm that the colored neighbors are genuine processes apposed to the spine within the tissue, not artifacts of the three-dimensional rendering, and show how the neuropil packs around the spine head at the nanometer scale. In this example, the final panel (H) adds a green glial process that occupies a large share of the neighborhood, enclosing several of the previously added elements.

**Figure 2.**
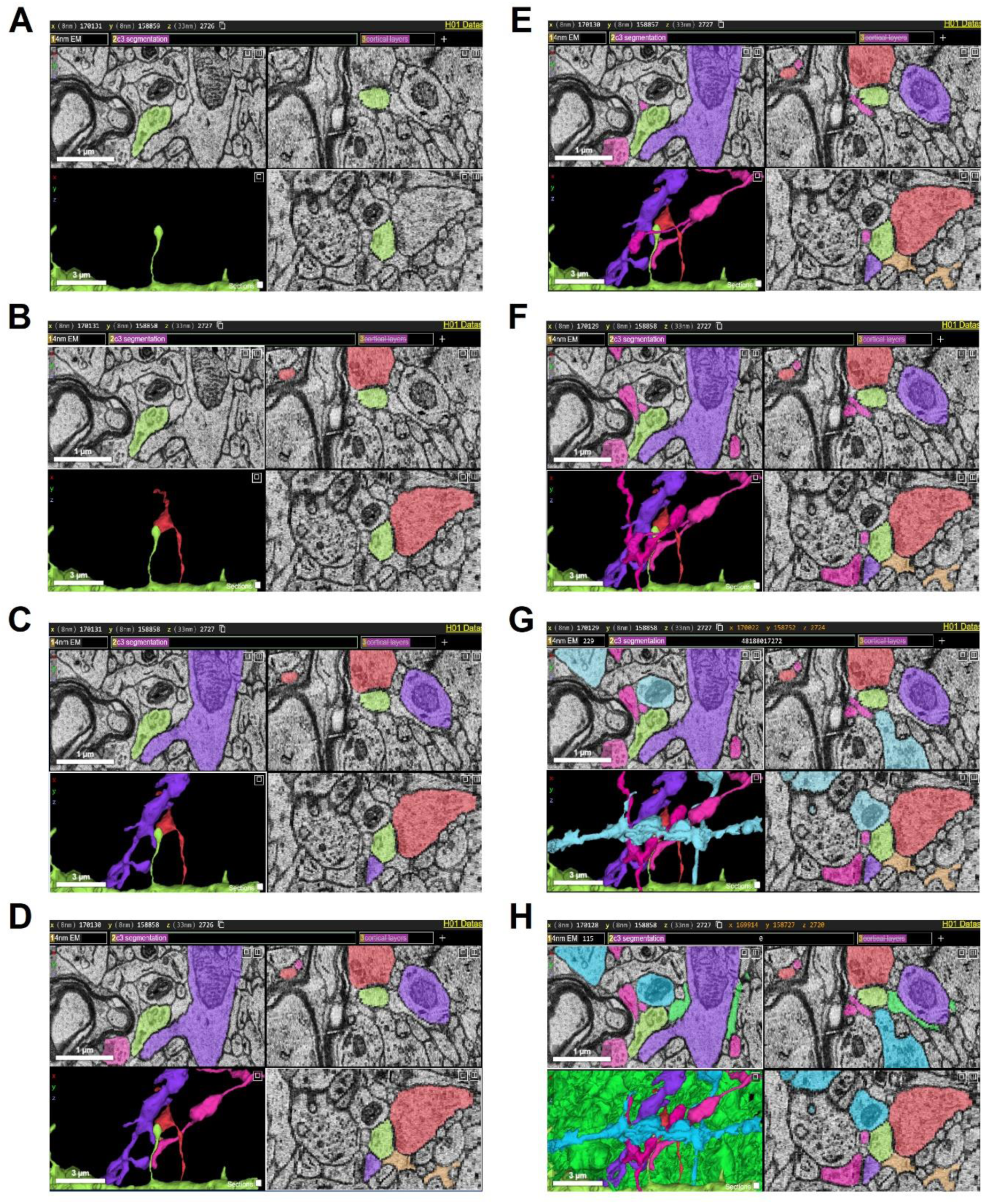
Neuronal neighborhood of a second layer 5 dendritic spine shown with orthogonal electron microscopy sections. A second dendritic spine (green) from the same layer 5 pyramidal neuron is shown with the surrounding neuronal elements, displayed as both three-dimensional renderings and corresponding electron microscopy sections through the spine head in orthogonal planes. **(A)** The spine alone. **(B through H)** The neighboring elements are added one at a time around the same spine, shown simultaneously in the three-dimensional reconstruction and in the cross-sectional views. The orthogonal sections confirm that the colored neighbors are genuine processes apposed to the spine within the tissue rather than artifacts of the three-dimensional rendering, and they show how densely the neuropil is packed around the spine head at the nanometer scale. Where two elements share a similar color, they correspond to separate processes from different neighboring neurons and should not be read as the same structure. Scale bars are indicated in each panel.

### Ground-truth analysis of the number of neighboring neuronal elements in the vicinome

After identifying each spine and its neighbors, we quantified two neighborhood properties across the full sample using ground-truth analysis (see Methods section). We characterized 4550 neuronal elements across the neuronal neighborhood of 322 dendritic spines from 17 pyramidal neurons across cortical layers 2 through 5, with each spine classified as apical or basal. For each spine, we quantified two properties over the same population of annotated vicinome neighbors. These were the number of neighboring neuronal elements and the minimum surface-to-surface distance between the spine and those elements. We defined groups by cortical layer crossed with spine polarity, yielding eight groups from L2 apical through L5 basal.

**Figure 3** illustrates how we measured the minimum distance on the ground-truth representations of a spine and one of its neighbors. From the twelve landmarks placed along each spine’s neck and head, through its reconstructed volume, to the smoothed mesh from which we took the measurement, the panels follow a single spine-neighbor pair to the point of closest approach, where the two surfaces lie 14 nm apart.

**Figure 3.**
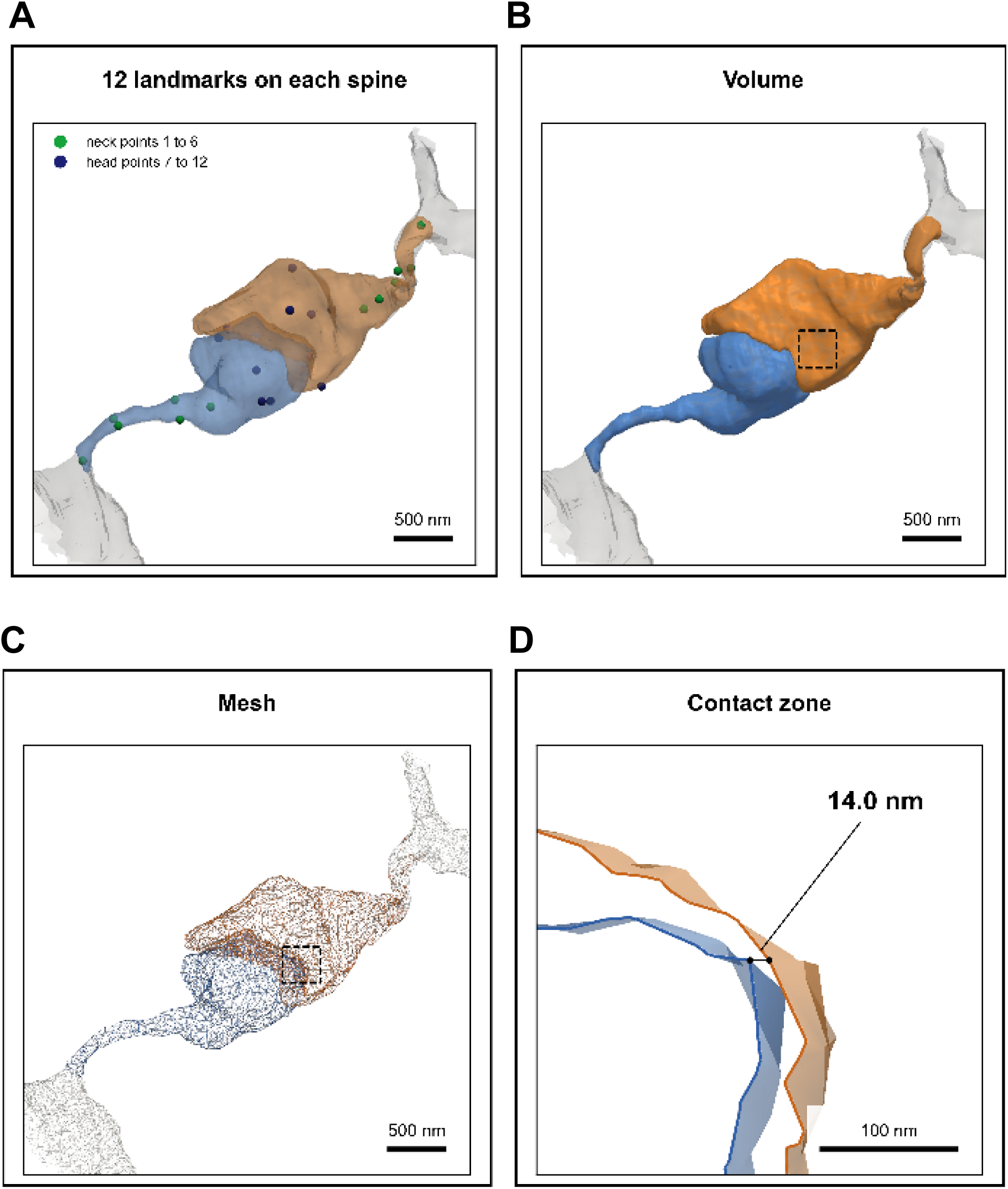
Measuring the minimum spine-to-neighbor distance on the ground-truth meshes. Four views of the same pair of neighboring elements, a spine (blue) and an adjacent process (orange), with the surrounding tissue in gray. **(A)** Twelve landmarks placed on the spine, six along the neck (green, points 1 to 6) and six on the head (dark blue, points 7 to 12). **(B)** The reconstructed volume of the two elements. **(C)** The smoothed surface mesh from which distances were computed. **(D)** The contact zone boxed in B and C, enlarged, where the two membranes reach their closest approach, and the minimum surface-to-surface distance was measured (14 nm). This pair comes from a single spine chosen at random to illustrate the procedure; its value is an individual example rather than representative of the group mean, which ranged from 30.4 to 34.6 nm across layers. Scale bars: 500 nm in A through C and 100 nm in D.

We found that the number of neighboring elements per spine decreased steadily from superficial to deep layers (**Figure 4**). Mean neighbor number fell from 17.2 ± 4.2 in L2 apical spines to 12.5 ± 3.1 in L5 basal spines. A Kruskal-Wallis test across the eight groups confirmed a global difference in neighboring number (chi-square 31.9, df 7, p = 4.3 x 10^-5^). Because spines are nested within neurons and are therefore not independent, we treated a linear mixed model with a random intercept per neuron as the primary analysis. The model showed a strong effect of cortical layer (F (3,314) = 7.59, p = 6.5 x 10^-5^) and no effect of apical versus basal polarity (F (1,314) = 0.24, p = 0.63), nor any layer-by-polarity interaction (F (3,314) = 0.32, p = 0.82). Relative to L2 apical spines, neighbor number decreased by 3.53 elements in L4 (p = 0.002) and by 3.83 in L5 (p = 0.001), whereas L3 did not differ significantly from L2 (p = 0.287).

**Figure 4.**
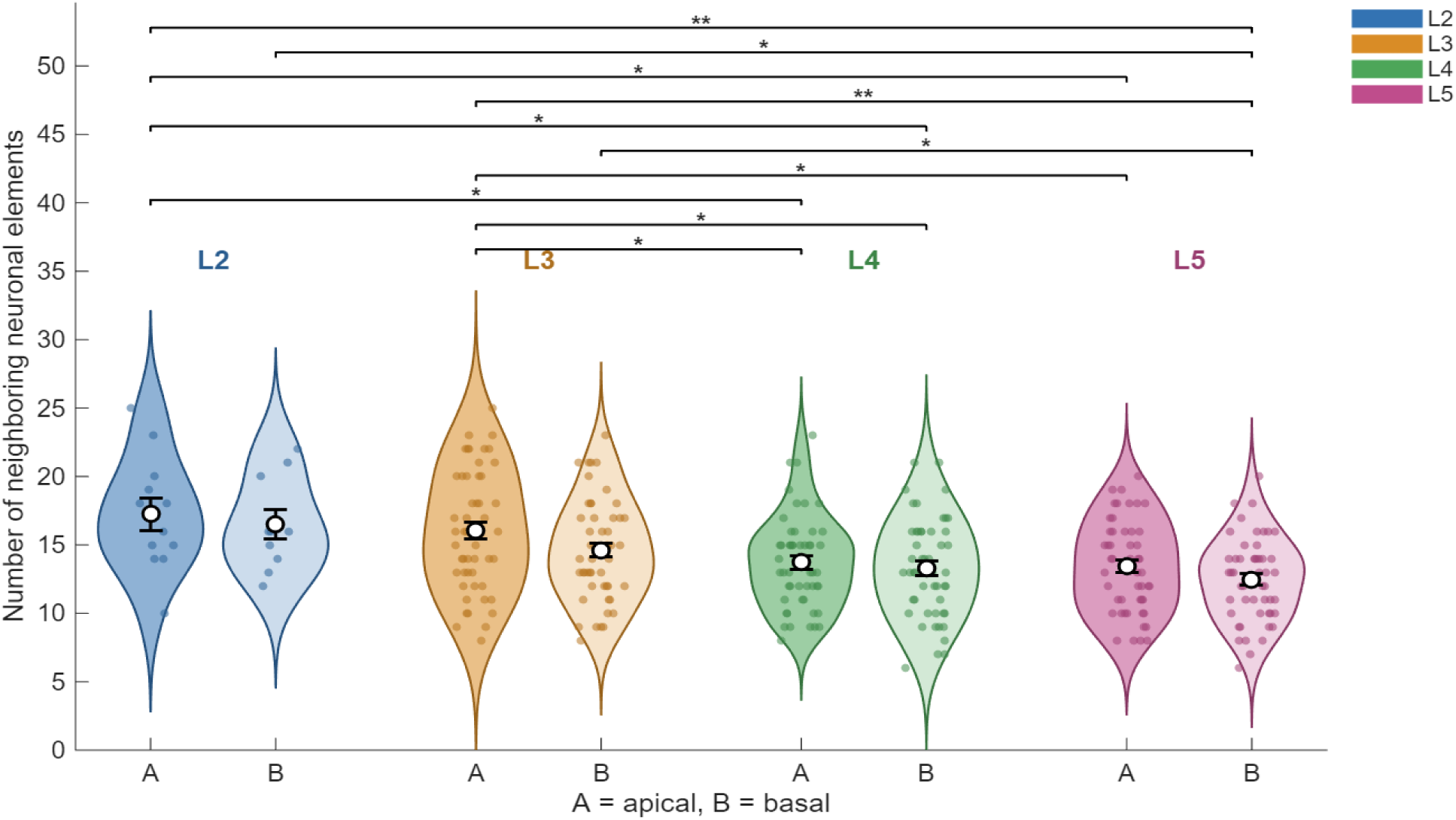
The number of neuronal elements neighboring a spine decreases with cortical depth. Violin plots of the number of neighboring neuronal elements per dendritic spine, for spines grouped by cortical layer (2 through 5) and by dendritic polarity (A, apical; B, basal). Each dot is one spine, the open circle marks the group mean, and the error bar shows the standard error of the mean. Neighbor number declines from superficial to deep layers, with no consistent difference between apical and basal spines. Brackets denote pairwise comparisons that remained significant after Benjamini Hochberg correction (Mann Whitney test); asterisks indicate the corrected significance level (one asterisk for p below 0.05, two for p below 0.01, three for p below 0.001). Group sizes, means, effect sizes, and full statistics are given in Table 1 (see also Table 2).

**Table 1.** Pairwise comparisons among the 8 groups (28 combinations) for the number of neighboring neuronal elements. Mann-Whitney test (two-sided) for each pair, with Benjamini-Hochberg FDR correction. Shown are the mean +/- SD of each group, the uncorrected p, the corrected p, and the Cohen’s d effect size (independent samples) with its 95% CI. * indicates corrected p < 0.05. ** indicates corrected p = 0.005 (Cohen d = 1.45), or p = 0.002 (Cohen d = 0.03) as indicated.

| Comparison | Mean +/- SD 1 | Mean +/- SD 2 | p (uncorrected) | p (FDR) | Cohen d | 95% CI of d | Sig. |
| --- | --- | --- | --- | --- | --- | --- | --- |
| L2A vs L2B | 17.2 +/- 4.2 | 16.5 +/- 3.4 | 0.715 | 0.792 | 0.20 | [-0.65, 1.04] |  |
| L2A vs L3A | 17.2 +/- 4.2 | 16.0 +/- 4.4 | 0.372 | 0.452 | 0.28 | [-0.35, 0.91] |  |
| L2A vs L3B | 17.2 +/- 4.2 | 14.6 +/- 3.7 | 0.048 | 0.096 | 0.70 | [0.05, 1.34] |  |
| L2A vs L4A | 17.2 +/- 4.2 | 13.7 +/- 3.4 | 0.008 | 0.025 | 0.99 | [0.34, 1.65] | * |
| L2A vs L4B | 17.2 +/- 4.2 | 13.3 +/- 3.6 | 0.005 | 0.022 | 1.06 | [0.40, 1.72] | * |
| L2A vs L5A | 17.2 +/- 4.2 | 13.4 +/- 3.4 | 0.006 | 0.022 | 1.09 | [0.43, 1.75] | * |
| L2A vs L5B | 17.2 +/- 4.2 | 12.5 +/- 3.1 | 0.0004 | 0.005 | 1.45 | [0.77, 2.13] | ** |
| L2B vs L3A | 16.5 +/- 3.4 | 16.0 +/- 4.4 | 0.735 | 0.792 | 0.11 | [-0.57, 0.79] |  |
| L2B vs L3B | 16.5 +/- 3.4 | 14.6 +/- 3.7 | 0.163 | 0.241 | 0.52 | [-0.17, 1.20] |  |
| L2B vs L4A | 16.5 +/- 3.4 | 13.7 +/- 3.4 | 0.025 | 0.054 | 0.82 | [0.12, 1.51] |  |
| L2B vs L4B | 16.5 +/- 3.4 | 13.3 +/- 3.6 | 0.024 | 0.054 | 0.89 | [0.19, 1.59] |  |
| L2B vs L5A | 16.5 +/- 3.4 | 13.4 +/- 3.4 | 0.023 | 0.054 | 0.92 | [0.22, 1.62] |  |
| L2B vs L5B | 16.5 +/- 3.4 | 12.5 +/- 3.1 | 0.002 | 0.021 | 1.29 | [0.57, 2.01] | * |
| L3A vs L3B | 16.0 +/- 4.4 | 14.6 +/- 3.7 | 0.125 | 0.206 | 0.34 | [-0.05, 0.74] |  |
| L3A vs L4A | 16.0 +/- 4.4 | 13.7 +/- 3.4 | 0.010 | 0.029 | 0.58 | [0.18, 0.98] | * |
| L3A vs L4B | 16.0 +/- 4.4 | 13.3 +/- 3.6 | 0.003 | 0.022 | 0.67 | [0.27, 1.07] | * |
| L3A vs L5A | 16.0 +/- 4.4 | 13.4 +/- 3.4 | 0.004 | 0.022 | 0.66 | [0.26, 1.06] | * |
| L3A vs L5B | 16.0 +/- 4.4 | 12.5 +/- 3.1 | 8.85 x 10<sup>-5</sup> | 0.002 | 0.93 | [0.52, 1.34] | ** |
| L3B vs L4A | 14.6 +/- 3.7 | 13.7 +/- 3.4 | 0.217 | 0.290 | 0.25 | [-0.14, 0.65] |  |
| L3B vs L4B | 14.6 +/- 3.7 | 13.3 +/- 3.6 | 0.112 | 0.195 | 0.36 | [-0.03, 0.76] |  |
| L3B vs L5A | 14.6 +/- 3.7 | 13.4 +/- 3.4 | 0.136 | 0.212 | 0.34 | [-0.05, 0.74] |  |
| L3B vs L5B | 14.6 +/- 3.7 | 12.5 +/- 3.1 | 0.006 | 0.022 | 0.63 | [0.23, 1.03] | * |
| L4A vs L4B | 13.7 +/- 3.4 | 13.3 +/- 3.6 | 0.724 | 0.792 | 0.12 | [-0.27, 0.51] |  |
| L4A vs L5A | 13.7 +/- 3.4 | 13.4 +/- 3.4 | 0.776 | 0.805 | 0.09 | [-0.30, 0.48] |  |
| L4A vs L5B | 13.7 +/- 3.4 | 12.5 +/- 3.1 | 0.098 | 0.183 | 0.38 | [-0.01, 0.78] |  |
| L4B vs L5A | 13.3 +/- 3.6 | 13.4 +/- 3.4 | 0.876 | 0.876 | -0.03 | [-0.43, 0.36] |  |
| L4B vs L5B | 13.3 +/- 3.6 | 12.5 +/- 3.1 | 0.265 | 0.337 | 0.24 | [-0.15, 0.64] |  |
| L5A vs L5B | 13.4 +/- 3.4 | 12.5 +/- 3.1 | 0.181 | 0.254 | 0.29 | [-0.10, 0.69] |  |

**Table 2.** Linear mixed model coefficients for the number of neighboring neuronal elements. Model: neighbors ∼ layer * polarity + (1|neuron). Reference: L2, apical. Random intercept per neuron. Reference = L2, apical. Each “L#” row is the difference of that layer relative to L2 (in apical spines). “basal” is the basal-apical difference in L2. The “L# x basal” rows are the interaction, that is, how much the basal-apical difference in that layer changes relative to L2.

| Term | beta (estimate) | SE | 95% CI of beta | t | df | p |
| --- | --- | --- | --- | --- | --- | --- |
| Intercept (L2, apical) | 17.250 | 1.036 | [15.211, 19.289] | 16.643 | 314 | 4.95 x 10<sup>-45</sup> |
| L3 | -1.230 | 1.154 | [-3.501, 1.041] | -1.066 | 314 | 0.287 |
| L4 | -3.530 | 1.154 | [-5.801, -1.259] | -3.059 | 314 | 0.002 |
| L5 | -3.830 | 1.154 | [-6.101, -1.559] | -3.319 | 314 | 0.001 |
| basal | -0.750 | 1.537 | [-3.775, 2.275] | -0.488 | 314 | 0.626 |
| L3 x basal | -0.650 | 1.697 | [-3.988, 2.688] | -0.383 | 314 | 0.702 |
| L4 x basal | 0.330 | 1.697 | [-3.008, 3.668] | 0.194 | 314 | 0.846 |
| L5 x basal | -0.190 | 1.697 | [-3.528, 3.148] | -0.112 | 314 | 0.911 |

Pairwise comparisons among the eight groups, corrected with the Benjamini-Hochberg procedure, placed every significant difference along the superficial-to-deep axis, and these effects were substantial in magnitude. The contrast between L2 apical and L5 basal spines reached a Cohen’s d of 1.45 (95% confidence interval 0.77 to 2.13), and L3 apical versus L5 basal reached a d of 0.93 (0.52 to 1.34), both large effects by conventional benchmarks.

Superficial groups also differed from deep ones at intermediate magnitudes; for example, L3 apical versus L4 basal showed d = 0.67 (0.27 to 1.07). No comparison between apical and basal spines within any layer was significant (**Table 1**). Cortical depth, and not dendritic polarity, thus governs how many neuronal elements surround a spine (**Table 1** and **Table 2**).

### Ground-truth analysis of the minimum spine-neighbor distance in the vicinome

In contrast to their number, the physical closeness of the neighbors in the vicinome was remarkably uniform across the cerebral cortex (**Figure 5**). The median minimum distance was near 28 nm in every group, and group means ranged narrowly from 30.4 to 34.6 nm, with a long right tail extending toward roughly 160 nm. A Kruskal-Wallis test detected a global difference across the eight groups (chi-square 42.4, df 7, p = 4.4 x 10^-7^), and several pairwise comparisons reached significance after correction, again distributed along the depth axis. However, these distance differences were negligible in magnitude. No significant contrast exceeded a Cohen’s d of 0.21, an effect below the conventional threshold for a small effect, and every confidence interval for d lay close to zero. The largest was L3 apical versus L5 basal, with a d of only 0.19 (95% confidence interval −0.29 to −0.08) despite a corrected p of 2.6 x 10^-6^. This illustrates how the large number of measured pairs confers statistical resolution on differences of a few nanometers that carry little biological weight.

**Figure 5.**
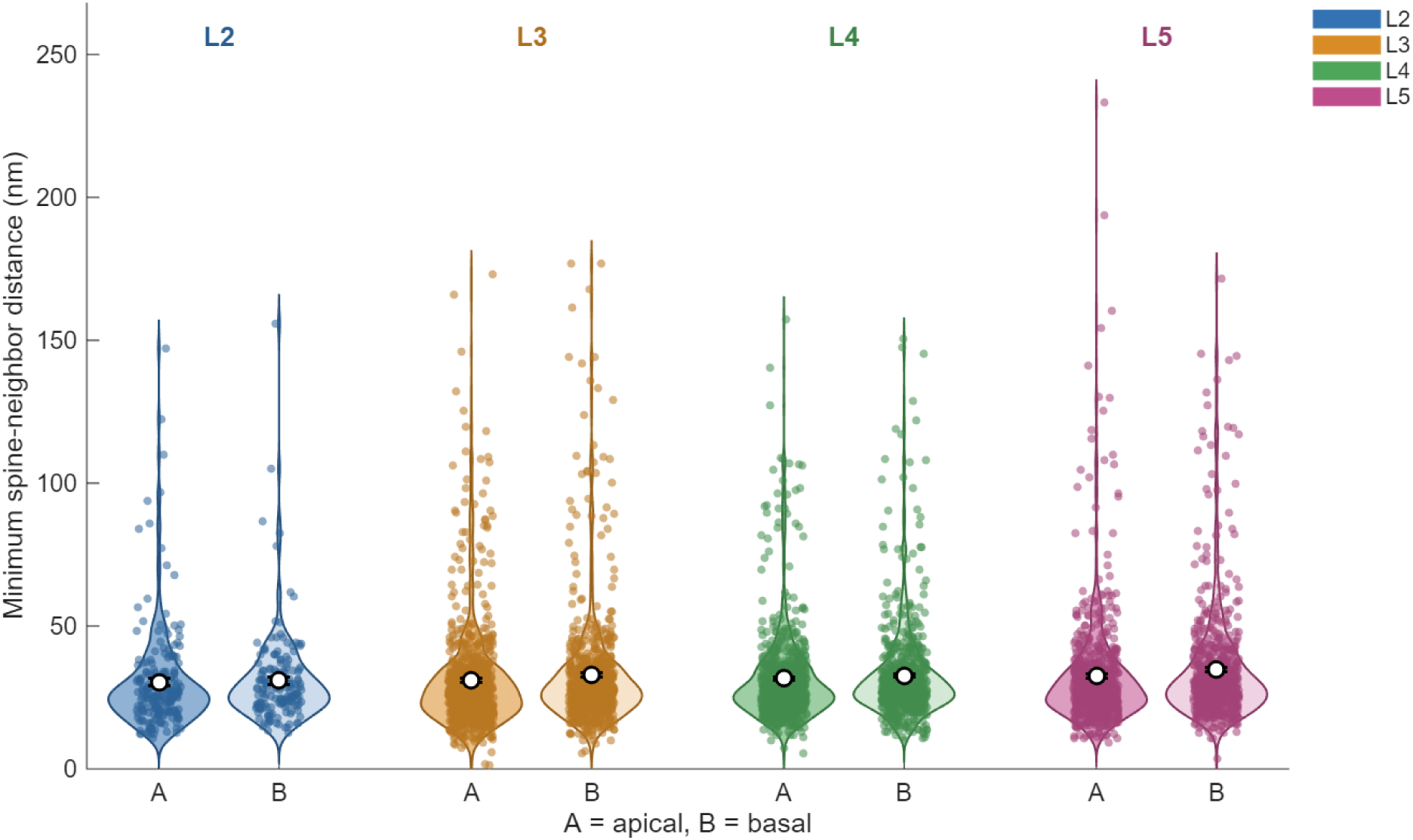
The minimum spine-to-neighbor distance is uniform across cortical layers and polarity. Violin plots of the minimum surface-to-surface distance (nm) between each dendritic spine and its neighboring neuronal elements, for spines grouped by cortical layer (2 through 5) and by dendritic polarity (A, apical; B, basal). Each dot is one spine-to-neighbor pair; the open circle marks the group means, and the error bar shows the standard error of the mean. The distributions are closely similar across all eight groups, with a median near 28 nm and a long tail toward larger distances. Pairwise comparisons are not marked on the plot because, although some reached statistical significance given the large number of pairs, all effect sizes were negligible (Cohen’s d below 0.21), so the differences carry little biological weight. Group sizes, means, corrected p-values, and effect sizes are reported in Table 3 (see also Table 4).

**Table 3.** Pairwise comparisons among the 8 groups (28 combinations) for the minimum spine-neighbor distance (nm). Mann-Whitney test (two-sided) for each pair, with Benjamini-Hochberg FDR correction. Shown are the mean +/- SD of each group, the uncorrected p, the corrected p, and the Cohen’s d effect size (independent samples) with its 95% CI. * indicates corrected p < 0.05; however, note that Cohen’s d effect sizes below 0.21 were negligible, which is why we read the minimum distance as essentially constant rather than as finely graded.

| Comparison | Mean +/- SD 1 | Mean +/- SD 2 | p (uncorrected) | p (FDR) | Cohen d | 95% CI of d | Sig. |
| --- | --- | --- | --- | --- | --- | --- | --- |
| L2A vs L2B | 30.4 +/- 18.4 | 30.9 +/- 16.3 | 0.152 | 0.218 | -0.03 | [-0.23, 0.18] |  |
| L2A vs L3A | 30.4 +/- 18.4 | 31.0 +/- 19.1 | 0.596 | 0.758 | -0.03 | [-0.18, 0.12] |  |
| L2A vs L3B | 30.4 +/- 18.4 | 32.8 +/- 21.1 | 0.020 | 0.047 | -0.11 | [-0.27, 0.04] | * |
| L2A vs L4A | 30.4 +/- 18.4 | 31.7 +/- 17.2 | 0.035 | 0.069 | -0.07 | [-0.23, 0.09] |  |
| L2A vs L4B | 30.4 +/- 18.4 | 32.6 +/- 17.5 | 0.0007 | 0.005 | -0.13 | [-0.28, 0.03] | * |
| L2A vs L5A | 30.4 +/- 18.4 | 32.4 +/- 20.6 | 0.030 | 0.064 | -0.10 | [-0.25, 0.06] |  |
| L2A vs L5B | 30.4 +/- 18.4 | 34.6 +/- 20.4 | 3.54 x 10<sup>-5</sup> | 0.0003 | -0.21 | [-0.37, -0.05] | * |
| L2B vs L3A | 30.9 +/- 16.3 | 31.0 +/- 19.1 | 0.226 | 0.302 | -0.00 | [-0.17, 0.16] |  |
| L2B vs L3B | 30.9 +/- 16.3 | 32.8 +/- 21.1 | 0.624 | 0.760 | -0.09 | [-0.26, 0.08] |  |
| L2B vs L4A | 30.9 +/- 16.3 | 31.7 +/- 17.2 | 0.790 | 0.850 | -0.04 | [-0.21, 0.13] |  |
| L2B vs L4B | 30.9 +/- 16.3 | 32.6 +/- 17.5 | 0.134 | 0.208 | -0.10 | [-0.27, 0.07] |  |
| L2B vs L5A | 30.9 +/- 16.3 | 32.4 +/- 20.6 | 0.728 | 0.815 | -0.08 | [-0.25, 0.09] |  |
| L2B vs L5B | 30.9 +/- 16.3 | 34.6 +/- 20.4 | 0.020 | 0.047 | -0.19 | [-0.36, -0.02] | * |
| L3A vs L3B | 31.0 +/- 19.1 | 32.8 +/- 21.1 | 0.007 | 0.025 | -0.09 | [-0.19, 0.01] | * |
| L3A vs L4A | 31.0 +/- 19.1 | 31.7 +/- 17.2 | 0.018 | 0.047 | -0.04 | [-0.14, 0.07] | * |
| L3A vs L4B | 31.0 +/- 19.1 | 32.6 +/- 17.5 | 3.54 x 10<sup>-5</sup> | 0.0003 | -0.09 | [-0.19, 0.01] | * |
| L3A vs L5A | 31.0 +/- 19.1 | 32.4 +/- 20.6 | 0.014 | 0.043 | -0.07 | [-0.17, 0.03] | * |
| L3A vs L5B | 31.0 +/- 19.1 | 34.6 +/- 20.4 | 9.13 x 10<sup>-8</sup> | 2.56 x 10<sup>-6</sup> | -0.19 | [-0.29, -0.08] | * |
| L3B vs L4A | 32.8 +/- 21.1 | 31.7 +/- 17.2 | 0.719 | 0.815 | 0.06 | [-0.05, 0.16] |  |
| L3B vs L4B | 32.8 +/- 21.1 | 32.6 +/- 17.5 | 0.133 | 0.208 | 0.01 | [-0.10, 0.11] |  |
| L3B vs L5A | 32.8 +/- 21.1 | 32.4 +/- 20.6 | 0.847 | 0.878 | 0.02 | [-0.09, 0.12] |  |
| L3B vs L5B | 32.8 +/- 21.1 | 34.6 +/- 20.4 | 0.004 | 0.015 | -0.09 | [-0.20, 0.02] | * |
| L4A vs L4B | 31.7 +/- 17.2 | 32.6 +/- 17.5 | 0.054 | 0.101 | -0.06 | [-0.16, 0.05] |  |
| L4A vs L5A | 31.7 +/- 17.2 | 32.4 +/- 20.6 | 0.888 | 0.888 | -0.04 | [-0.15, 0.07] |  |
| L4A vs L5B | 31.7 +/- 17.2 | 34.6 +/- 20.4 | 0.001 | 0.006 | -0.16 | [-0.27, -0.05] | * |
| L4B vs L5A | 32.6 +/- 17.5 | 32.4 +/- 20.6 | 0.081 | 0.141 | 0.01 | [-0.09, 0.12] |  |
| L4B vs L5B | 32.6 +/- 17.5 | 34.6 +/- 20.4 | 0.155 | 0.218 | -0.11 | [-0.21, 0.00] |  |
| L5A vs L5B | 32.4 +/- 20.6 | 34.6 +/- 20.4 | 0.002 | 0.012 | -0.11 | [-0.22, -0.00] | * |

**Table 4.** Linear mixed model coefficients for the minimum spine-neighbor distance (nm). Model: neighbors ∼ layer * polarity + (1|neuron). Reference: L2, apical. Random intercept per neuron. Reference = L2, apical. Each “L#” row is the difference of that layer relative to L2 (in apical spines). “basal” is the basal-apical difference in L2. The “L# x basal” rows are the interaction, that is, how much the basal-apical difference in that layer changes relative to L2.

| Term | beta (estimate) | SE | 95% CI of beta | t | df | p |
| --- | --- | --- | --- | --- | --- | --- |
| Intercept (L2, apical) | 30.423 | 1.335 | [27.806, 33.040] | 22.793 | 4542 | 5.65 x 10<sup>-109</sup> |
| L3 | 0.554 | 1.497 | [-2.382, 3.489] | 0.370 | 4542 | 0.712 |
| L4 | 1.228 | 1.523 | [-1.758, 4.214] | 0.806 | 4542 | 0.420 |
| L5 | 1.970 | 1.527 | [-1.023, 4.964] | 1.290 | 4542 | 0.197 |
| basal | 0.471 | 2.004 | [-3.458, 4.400] | 0.235 | 4542 | 0.814 |
| L3 x basal | 1.320 | 2.232 | [-3.055, 5.696] | 0.592 | 4542 | 0.554 |
| L4 x basal | 0.524 | 2.260 | [-3.907, 4.955] | 0.232 | 4542 | 0.817 |
| L5 x basal | 1.782 | 2.271 | [-2.670, 6.235] | 0.785 | 4542 | 0.433 |

Consistent with this, the linear mixed model found no significant effect of layer on distance once neuron-level nesting was accounted for (F (3,4542) = 0.91, p = 0.43), and no effect of polarity (F (1,4542) = 0.06, p = 0.81). We therefore regard the minimum spine-to-neighbor distance as essentially constant across layers and polarity, near 30 nm, a value close to the scale of the chemical synaptic cleft and compatible with chemical, electrical, or potentially ephaptic apposition, although electron microscopy alone cannot establish functional interaction (**Table 3** and **Table 4**).

## Discussion

Taken together, the two measurements describe a dendritic spine vicinome whose composition changes with cortical depth while its geometry does not. Deeper spines are surrounded by fewer neuronal elements, a graded and sizable effect, yet those elements sit just as close to the spine surface as in the superficial layers, where neighbors are more numerous. Dendritic polarity influenced neither property.

The falling neighbor count within the vicinome with depth invites comparison with what volume electron microscopy has reported about how densely the neuropil is packed across layers. Santuy et al. (2018) reconstructed 6184 synaptic junctions through the six layers of the juvenile rat somatosensory cortex and found that synaptic density itself varies by layer, highest in layers II and IV and lowest in layer VI, with excitatory contacts preferring spines and inhibitory ones the shafts. Our measurement is not of synapses but of vicinome neighboring elements, and our tissue is adult human temporal cortex; hence, we cannot equate the two. Still, it is interesting that both descriptions place the superficial layers as more crowded and the deep layers as sparser.

That a single spine can be flanked by many elements also recalls how many contacts a spine can itself receive. Hwang et al. (2021) showed with serial block face imaging that a spine on a GABAergic interneuron in mouse visual cortex can carry as many as eight synaptic inputs, and Bosch et al. (2015), using FIB/SEM in labeled adult-generated granule cells, found presynaptic terminals contacting up to ten dendritic spines. Read alongside our counts of twelve to seventeen neighboring elements per spine, these findings sketch a picture in which neither the spine nor the bouton behaves as an isolated endpoint. We counted elements in physical proximity in the vicinome rather than confirmed synapses, so our numbers and theirs measure different things, and we draw only the modest inference that the immediate surround of a spine is populated rather than empty.

The uniformity of the minimum distance, near 30 nm, sits close to the scale at which synaptic ultrastructure is ordinarily described. Harris and Weinberg (2012) reviewed the fine structure of excitatory synapses and placed the synaptic cleft at a few tens of nanometers, with the postsynaptic density marking the spine as the canonical glutamatergic target and inhibitory contacts settling on shafts and somata. Our vicinome group means, ranging narrowly from 30.4 to 34.6 nm with a median near 28 nm, fall within that same range, though we measured the nearest approach of any neighboring element rather than synaptic apposition. The coincidence of scale suggests a floor on how close membranes lie and we leave it at that, since anatomy alone cannot tell whether a given apposition is a synapse.

Our reliance on the ground-truth meshes rather than on projected visual estimates rests on the resolving power that volume techniques now afford. Takahashi-Nakazato et al. (2019) documented that FIB/SEM, with *en bloc* heavy metal staining, resolves postsynaptic densities and plasma membrane contours cleanly enough to measure the features that shape glutamatergic transmission, and Parajuli and Koike (2021) reviewed how volumetric imaging has become the standard for the precise quantitation of synaptic parameters. These accounts concern the accuracy of the method rather than any biological claim of ours, and we cite them to support the plausibility of surface-to-surface distances measured in the tens of nanometers, a precision beyond the reach of projected two-dimensional estimates.

Researchers have previously reported a geometric regularity in how spines are arranged along the dendrite rather than across the layers. Parajuli et al. (2020) found in several mouse regions that the postsynaptic density area per unit length of dendrite scales with dendritic diameter, and that the ratio of postsynaptic density area to neck length stays relatively uniform, an organization they linked through simulation to a comparable synaptic strength across dendrites. Our finding concerns a different type of variation. The number of neighbors in the dendritic spine vicinome falls with cortical depth while their proximity remains constant, and we do not know whether the two regularities share any mechanism. The comparison points to a common theme: that the spine and its surroundings in the vicinome appear organized rather than random, which in our data takes the form of one property that varies with depth and one that holds constant.

Reading the spine within its vicinome, rather than as a point in isolation, is the reframing that Kuwajima et al. (2013) urged for studying spine pathology, arguing that number and shape acquire meaning only when the presynaptic partners, the astrocytic processes, and the surrounding organelles are seen as well. Our characterization applies that reframing to the healthy human spine, quantifying the surround as an object and treating it as the object of measurement. We describe the surround anatomically as a first step, without making functional claims, and with the caution the approach itself imposes.

Among those surrounding elements in the vicinome, glial processes deserve particular mention. Rollenhagen et al. (2025), reconstructing excitatory boutons in layer 1 of the human temporal neocortex, described astrocytic coverage of synaptic complexes consistent with both synaptic crosstalk and the removal of spilled glutamate. In our sample, glial processes recurred among the neighbors and at times occupied a large share of the neighborhood, as the reconstruction in Figure 2H illustrates. We identified these processes by proximity and did not assess their function, so the correspondence is anatomical, a reminder that the neighborhood of a human spine includes non-neuronal members whose closeness may matter.

Placing this work in the small body of human volume electron microscopy locates both its footing and its limits. Glausier et al. (2025) recovered intact ultrastructural correlates of synaptic function in postmortem human prefrontal cortex; Cano-Astorga et al. (2021) and Rollenhagen and Lübke (2025) quantified the synaptic organization of the human temporal neocortex from autopsy and biopsy tissue; and Domínguez-Álvaro et al. (2019) extended such analysis to the transentorhinal cortex, where the proportion of synapses on spine heads was reduced in Alzheimer’s disease. Our study shares their substrate, human temporal cortex, and their premise that three-dimensional ultrastructure is now quantifiable in human tissue, while asking a question they did not: how many elements encircle a spine and how near they lie within the vicinome. The pathological contrast that Domínguez-Álvaro et al. (2019) drew suggests that a baseline description of the healthy neighborhood, such as ours, could one day serve as a reference against which altered states are compared, a possibility we raise without testing it here.

If any of this matters functionally, it is due to the proximity itself. The extracellular fields generated by active neurons can influence nearby membranes independently of chemical or electrical synapses, an ephaptic coupling strongest where elements lie close together and at slow timescales (Anastassiou et al., 2011; Han et al., 2018), and the spilled glutamate that astrocytes recover can reach receptors on membranes that form no synapse (Rollenhagen et al., 2025; Kwon et al., 2019). A minimum distance near 30 nm across all layers is the anatomical condition such interactions would require, though it is a necessary condition and not evidence that they occur. Electron microscopy fixes the geometry and leaves physiology open, and we interpret our findings cautiously within that boundary.

## Limitations

Our study has several limitations. The reconstruction derives from a single donor, so the depth gradient we report awaits confirmation across individuals. Neighbors were identified by manual expert inspection before measurement on the meshes, a step that carries the observer’s judgment even when the subsequent distances are objective. The number of measured pairs is large enough that trivial distance differences reached statistical significance while carrying negligible effect sizes, all below a Cohen’s d of 0.21, which is why we read the minimum distance as essentially constant rather than as finely graded. And proximity, throughout, is an anatomical measure. Whether the crowding we describe shapes how a spine integrates its inputs is an open question that this method poses but cannot answer.

### Perspectives and future directions

Our study characterizes the dendritic spine as a structure embedded in a vicinome with crowded surroundings, which, once described, invites the question of what it does. Physiology has long been understood as structure in action, in the spirit of Sherrington’s integrative view of the nervous system (Sherrington, 1906), and the vicinome crowding we measured is, for now, only the structure. What that vicinome crowding does to the spine, whether it shapes how the spine integrates its inputs, remains outside what electron microscopy can reach. In the next paragraphs, we raise several possibilities as directions for research, not as findings, mindful that anatomy fixes the geometry and leaves physiology open.

The proximity we report within the vicinome, a minimum distance near 30 nm holding across every layer, is the anatomical condition that several non-classical influences would require. Extracellular fields from active neurons can act on nearby membranes through ephaptic coupling, strongest where elements lie close and at slow timescales (Anastassiou et al., 2011; Han et al., 2018), and glutamate that escapes the cleft can reach extrasynaptic receptors on membranes that form no synapse (Kwon et al., 2019; Rollenhagen et al., 2025). A spine encircled by many tightly apposed neighbors within the vicinome therefore sits in a setting where such influences are geometrically possible. Whether they operate, and whether their strength tracks the number of neighbors we counted, is a question our data pose but cannot settle. We state plainly that ephaptic or extrasynaptic action cannot be inferred from anatomy alone.

Yet the inability to infer function from structure is not a reason to leave the question unasked. Science advances in part by proposing fields of study before the tools to close them exist, and a structural description like ours can mark where those tools should be aimed. We suggest that the spine’s crowded neighborhood in the vicinome is worth modeling and testing directly. Computational models that place the measured neighbors, at their measured distances, around a reconstructed spine could estimate whether the extracellular fields of active neighbors perturb the spine’s membrane potential, and whether that perturbation scales with neighbor number as it falls from the superficial to the deep layers.

It is tempting to speculate that experiments combining spine-resolution imaging with manipulation of nearby activity could ask whether a spine’s excitability shifts with the activity of elements that never contact it across a synapse. The ground-truth meshes we used supply the geometry such models require, and the natural next step is to pair them with physiology.

The characterization could also be widened. We identified neighbors in the vicinome by manual expert inspection, a reliable but slow procedure that limited the analysis to 322 spines from a single donor. Automated segmentation by deep learning, already under development for volumes of this kind, would scale the vicinome analysis to thousands of spines and several donors, and that reach would bring the statistical power to separate effects by cell type, not only by layer and polarity. What we present as a depth gradient in one brain could then be tested for its generality and its finer structure.

We also raise a translational horizon, with caution. Since altered targeting of dendritic spines has been reported in disease, with a reduced proportion of synapses on spine heads in Alzheimer’s disease (Domínguez-Álvaro et al., 2019), or altered synaptic plasticity (Zhang et al., 2022) with lower densities of spines in schizophrenia (Fish et al., 2025), a baseline description of the healthy spine vicinome could one day serve as a reference against which pathological states are examined. We could ask whether the crowding or the proximity in the vicinome we measured shifts when the circuit is diseased. We name this only as a horizon, since our sample is a single healthy donor and we have tested no clinical contrast.

It is tempting to speculate that crowding of elements around a dendritic spine is an observable that could change with disease. The two properties we measured, the number of neighbors and their minimum distance, are one way to render that observable concrete, and a measure of vicinome crowding built from them could travel across brain regions, ages, species, and clinical conditions. We offer the term as an invitation to measurement and not as a claim, since whether vicinome crowding carries functional or clinical importance is exactly what future work would have to establish.

Framed this way, our characterization serves as a substrate for a broader program: studying how ephaptic and extrasynaptic influences from many neighbors contribute to dendritic spine function. We do not claim that program’s conclusions in advance. We propose that the vicinome crowding is real, that it is graded by cortical depth, that its nearest members lie within reach of non-synaptic influence, and that these facts, taken together, make the functional question worth pursuing.

The spine has been studied for a century as a point. Seen as a hub in a dense vicinome, it becomes a place where structure and its possible actions can be asked about together. By giving the neighborhood a name (the vicinome) and a measure (the degree of crowding), we hope to open a line of inquiry that anatomy alone cannot close, one that asks how the crowding of the vicinome shapes the physiology of the spine and, further still, whether that crowding differs in the diseased brain. A disorder that reorganizes the brain might leave its trace in a denser or sparser dendritic spine vicinome, and a measurable trace is the first step toward a mechanism. Here we take only the first step: the anatomy, describing the healthy vicinome and its crowding in a single human brain. Physiology and the comparison across health and disease are the steps that would follow. If the vicinome varies in meaningful ways, neuroscience will have gained not just a word but a window.

## Conclusions

The human dendritic spine could operate within a crowded envelope of neighboring elements whose density is set by laminar position, but whose nearest members remain within a few tens of nanometers regardless of layer. We name this envelope the dendritic spine vicinome and propose its crowding (denser or sparser) as a quantifiable observable, one whose functional and clinical relevance future work will have to test.

## Data Availability Statement

Source data for this study will be openly available in a repository after acceptance.

## Author contributions

EM conceived and designed the study and wrote the paper. All authors analyzed the database, revised the manuscript, and approved it.

## Ethical Approval Statement

This manuscript does not require an ethical approval statement, as it was performed on a publicly available database.

## Acknowledgments

NL acknowledges a Master of Science fellowship from SECIHTI, and MAZU acknowledges a postdoctoral fellowship from SECIHTI. AF and EM acknowledge support from VIEP-BUAP and Fundación Marcos Moshinsky.

## References

Anastassiou, C. A., Perin, R., Markram, H., & Koch, C. (2011). Ephaptic coupling of cortical neurons. Nature neuroscience, 14(2), 217–223. 10.1038/nn.2727

Arellano, J. I., Benavides-Piccione, R., Defelipe, J., & Yuste, R. (2007). Ultrastructure of dendritic spines: correlation between synaptic and spine morphologies. Frontiers in neuroscience, 1(1), 131–143. 10.3389/neuro.01.1.1.010.2007

Benavides-Piccione R, Fernaud-Espinosa I, Robles V, Yuste R, DeFelipe J. (2013) Age-based comparison of human dendritic spine structure using complete three-dimensional reconstructions. Cereb Cortex. 23(8), 1798–810. doi: 10.1093/cercor/bhs154. Epub 2012 Jun 17. PMID: 22710613; PMCID: PMC3698364.

Bosch, C., Martínez, A., Masachs, N., Teixeira, C. M., Fernaud, I., Ulloa, F., Pérez-Martínez, E., Lois, C., Comella, J. X., DeFelipe, J., Merchán-Pérez, A., & Soriano, E. (2015). FIB/SEM technology and high-throughput 3D reconstruction of dendritic spines and synapses in GFP-labeled adult-generated neurons. Frontiers in neuroanatomy, 9, 60. 10.3389/fnana.2015.00060

Bourne, J. N., & Harris, K. M. (2008). Balancing structure and function at hippocampal dendritic spines. Annual review of neuroscience, 31, 47–67. 10.1146/annurev.neuro.31.060407.125646

Caire, M. J., Reddy, V., & Varacallo, M. A. (2023). Physiology, Synapse. In StatPearls. StatPearls Publishing.

Cano-Astorga, N., DeFelipe, J., & Alonso-Nanclares, L. (2021). Three-Dimensional Synaptic Organization of Layer III of the Human Temporal Neocortex. Cerebral cortex (New York, N.Y.: 1991), 31(10), 4742–4764. 10.1093/cercor/bhab120

Domínguez-Álvaro, M., Montero-Crespo, M., Blazquez-Llorca, L., DeFelipe, J., & Alonso-Nanclares, L. (2019). 3D Electron Microscopy Study of Synaptic Organization of the Normal Human Transentorhinal Cortex and Its Possible Alterations in Alzheimer’s Disease. eNeuro, 6(4), ENEURO.0140-19.2019. 10.1523/ENEURO.0140-19.2019

Fish KN, Sweet RA, MacDonald ML, Lewis DA. Regional Specificity of Cortical Layer 3 Dendritic Spine Deficits in Schizophrenia. JAMA Psychiatry. 2025 Nov 1;82(11):1123–1132. doi: 10.1001/jamapsychiatry.2025.2221. PMID: 40960807; PMCID: PMC12444653.

García-López, P., García-Marín, V., & Freire, M. (2007). The discovery of dendritic spines by Cajal in 1888 and its relevance in the present neuroscience. Progress in neurobiology, 83(2), 110–130. 10.1016/j.pneurobio.2007.06.002.

García-López, P., García-Marín, V., & Freire, M. (2010). Dendritic spines and development: towards a unifying model of spinogenesis--a present day review of Cajal’s histological slides and drawings. Neural plasticity, 2010, 769207. 10.1155/2010/769207

Geckt, D., Ofer, N., Reimann, M.W., Yuste, R., Segev, I. (2026) Dendritic spines implement specific connectivity and support neuronal ensembles in cortical circuits. bioRxiv [Preprint]. 2026 Aug 17:2026.06.07.730704. doi: 10.64898/2026.06.07.730704. PMID: 42327032; PMCID: PMC13277910.

Glausier, J. R., Maier, M., Bouchet-Marquis, C., Wu, K., Banks-Tibbs, T., Melchitzky, D., Ning, J., Lewis, D. A., & Freyberg, Z. (2025). Volume electron microscopy reveals 3D synaptic nanoarchitecture in postmortem human prefrontal cortex. iScience, 28(7), 112747. 10.1016/j.isci.2025.112747

Han, K. S., Guo, C., Chen, C. H., Witter, L., Osorno, T., & Regehr, W. G. (2018). Ephaptic Coupling Promotes Synchronous Firing of Cerebellar Purkinje Cells. Neuron, 100(3), 564–578.e3. 10.1016/j.neuron.2018.09.018

Harris KM, Stevens JK. (1989). Dendritic spines of CA 1 pyramidal cells in the rat hippocampus: serial electron microscopy with reference to their biophysical characteristics. J Neurosci. 9(8),2982–97. doi: 10.1523/JNEUROSCI.09-08-02982.1989.

Harris, K. M., & Weinberg, R. J. (2012). Ultrastructure of synapses in the mammalian brain. Cold Spring Harbor perspectives in biology, 4(5), a005587. 10.1101/cshperspect.a005587

Hwang, Y. S., Maclachlan, C., Blanc, J., Dubois, A., Petersen, C. C. H., Knott, G., & Lee, S. H. (2021). 3D Ultrastructure of Synaptic Inputs to Distinct GABAergic Neurons in the Mouse Primary Visual Cortex. Cerebral cortex (New York, N.Y.: 1991), 31(5), 2610–2624. 10.1093/cercor/bhaa378

Januszewski M, Kornfeld J, Li PH, Pope A, Blakely T, Lindsey L, Maitin-Shepard J, Tyka M, Denk W, Jain V. High-precision automated reconstruction of neurons with flood-filling networks. Nature Methods. 2018;15(8):605–610. doi:10.1038/s41592-018-0049-4. PMID: 30013046.

Kubota, Y., Sohn, J., & Kawaguchi, Y. (2018). Large Volume Electron Microscopy and Neural Microcircuit Analysis. Frontiers in neural circuits, 12, 98. 10.3389/fncir.2018.00098

Kuwajima, M., Spacek, J., & Harris, K. M. (2013). Beyond counts and shapes: studying pathology of dendritic spines in the context of the surrounding neuropil through serial section electron microscopy. Neuroscience, 251, 75–89. 10.1016/j.neuroscience.2012.04.061

Kwon, T., Merchán-Pérez, A., Rial Verde, E. M., Rodríguez, J. R., DeFelipe, J., & Yuste, R. (2019). Ultrastructural, Molecular and Functional Mapping of GABAergic Synapses on Dendritic Spines and Shafts of Neocortical Pyramidal Neurons. Cerebral cortex (New York, N.Y.: 1991), 29(7), 2771–2781. 10.1093/cercor/bhy143

Manjarrez, E., Hernandez, S.T., Zamora-Ursulo, M.A., Flores, A. (2026) Rotating a petavoxel reconstruction exposes the viewing-angle bias inherent to Golgi-Cox and confocal dendritic-spine classification. Journal of Neurophysiology. In press.

Ofer, N., Benavides-Piccione, R., DeFelipe, J., & Yuste, R. (2022). Structural Analysis of Human and Mouse Dendritic Spines Reveals a Morphological Continuum and Differences across Ages and Species. eNeuro, 9(3), ENEURO.0039-22.2022. 10.1523/ENEURO.0039-22.2022

Parajuli, L. K., & Koike, M. (2021). Three-Dimensional Structure of Dendritic Spines Revealed by Volume Electron Microscopy Techniques. Frontiers in neuroanatomy, 15, 627368. 10.3389/fnana.2021.627368

Parajuli, L. K., Urakubo, H., Takahashi-Nakazato, A., Ogelman, R., Iwasaki, H., Koike, M., Kwon, H. B., Ishii, S., Oh, W. C., Fukazawa, Y., & Okabe, S. (2020). Geometry and the Organizational Principle of Spine Synapses along a Dendrite. eNeuro, 7(6), ENEURO.0248-20.2020. 10.1523/ENEURO.0248-20.2020

Reberger, R., Dall’Oglio, A., Jung, C. R., & Rasia-Filho, A. A. (2018). Structure and diversity of human dendritic spines evidenced by a new three-dimensional reconstruction procedure for Golgi staining and light microscopy. Journal of neuroscience methods, 293, 27–36. 10.1016/j.jneumeth.2017.09.001

Renner, J., & Rasia-Filho, A. A. (2023). Morphological Features of Human Dendritic Spines. Advances in neurobiology, 34, 367–496. 10.1007/978-3-031-36159-3_9

Rochefort, N. L., & Konnerth, A. (2012). Dendritic spines: from structure to in vivo function. EMBO reports, 13(8), 699–708. 10.1038/embor.2012.102

Rollenhagen, A., & Lübke, J. H. R. (2025). The synaptic organization of the human temporal lobe neocortex by high-resolution transmission, focused ion beam scanning, and electron microscopic tomography. Anatomical science international, 100(4), 480–497. 10.1007/s12565-025-00900-y

Rollenhagen, A., Sadeghi, A., Walkenfort, B., Hilgetag, C. C., Sätzler, K., & Lübke, J. H. R. (2025). Ultrastructural sublaminar-specific diversity of excitatory synaptic boutons in layer 1 of the adult human temporal lobe neocortex. eLife, 13, RP99473. 10.7554/eLife.99473

Santuy, A., Rodriguez, J. R., DeFelipe, J., & Merchan-Perez, A. (2018). Volume electron microscopy of the distribution of synapses in the neuropil of the juvenile rat somatosensory cortex. Brain structure & function, 223(1), 77–90. 10.1007/s00429-017-1470-7

Shapson-Coe A, Januszewski M, Berger DR, Pope A, Wu Y, Blakely T, Schalek RL, Li PH, Wang S, Maitin-Shepard J, Karlupia N, Dorkenwald S, Sjostedt E, Leavitt L, Lee D, Troidl J, Collman F, Bailey L, Fitzmaurice A, Kar R, Field B, Wu H, Wagner-Carena J, Aley D, Lau J, Lin Z, Wei D, Pfister H, Peleg A, Jain V, Lichtman JW. (2024). A petavoxel fragment of human cerebral cortex reconstructed at nanoscale resolution. Science (New York, N.Y.), 384(6696), eadk4858. 10.1126/science.adk4858

Sherrington, C. S. (1906). The Integrative Action of the Nervous System. New York: Charles Scribner’s Sons.

Südhof T. C. (2018). Towards an Understanding of Synapse Formation. Neuron, 100(2), 276–293. 10.1016/j.neuron.2018.09.040

Takahashi-Nakazato, A., Parajuli, L. K., Iwasaki, H., Tanaka, S., & Okabe, S. (2019). Ultrastructural Observation of Glutamatergic Synapses by Focused Ion Beam Scanning Electron Microscopy (FIB/SEM). Methods in molecular biology (Clifton, N.J.), 1941, 17–27. 10.1007/978-1-4939-9077-1_2

Tønnesen, J., & Nägerl, U. V. (2016). Dendritic Spines as Tunable Regulators of Synaptic Signals. Frontiers in psychiatry, 7, 101. 10.3389/fpsyt.2016.00101

Wilson, C. J., Groves, P. M., Kitai, S. T., & Linder, J. C. (1983). Three-dimensional structure of dendritic spines in the rat neostriatum. The Journal of neuroscience: the official journal of the Society for Neuroscience, 3(2), 383–388. 10.1523/JNEUROSCI.03-02-00383.1983

Yuste R. (2015). The discovery of dendritic spines by Cajal. Front Neuroanat. 2015 Apr 21;9:18. doi: 10.3389/fnana.2015.00018. PMID: 25954162; PMCID: PMC4404913.

Zamora-Ursulo M.A. & Manjarrez, E. (2026). Orientation-invariant morphometry reveals a continuum of dendritic spine forms in layer II pyramidal neurons of the petavoxel human connectome. bioRxiv. 10.64898/2026.06.25.734571

Zhang K, Liao P, Wen J, Hu Z. Synaptic plasticity in schizophrenia pathophysiology. IBRO Neurosci Rep. 2022 Oct 31;13:478–487. doi: 10.1016/j.ibneur.2022.10.008. PMID: 36590092; PMCID: PMC9795311.

